## Supplementary figures and images for "Astrocytes modulate neuronal development by S100A6 signaling"

### Supplementary Figure 1

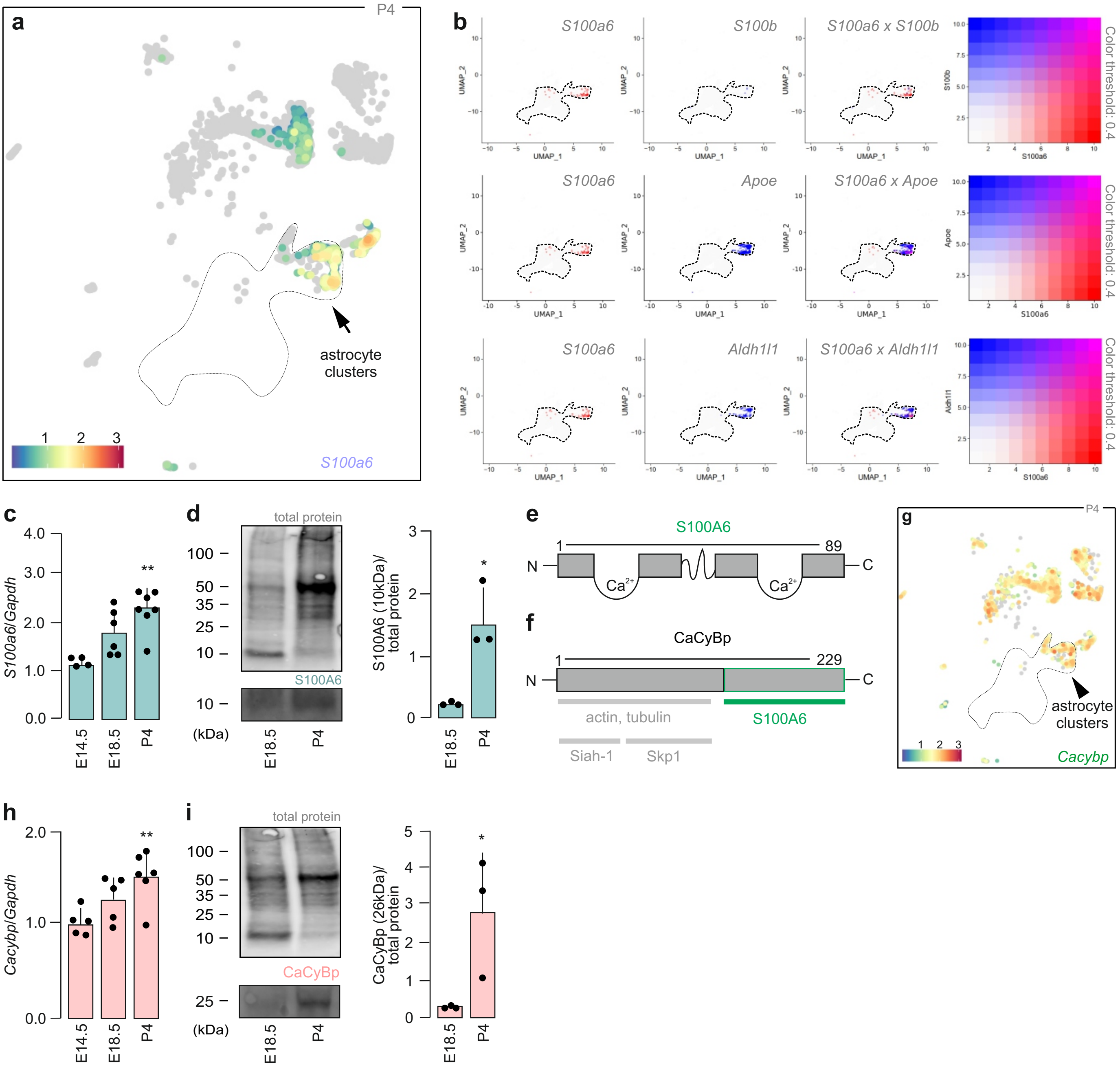

### Supplementary Figure 2

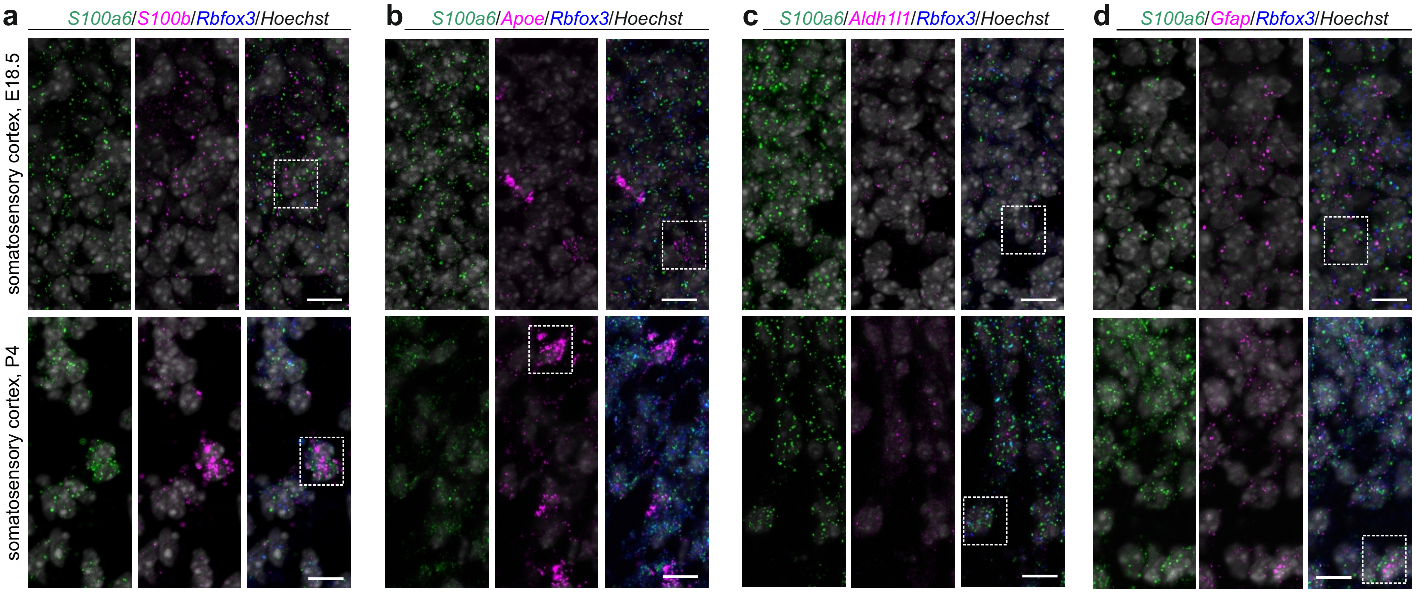

### Supplementary Figure 3

Cinquina *et al*. - Supplementary Fig. 3 (revision)

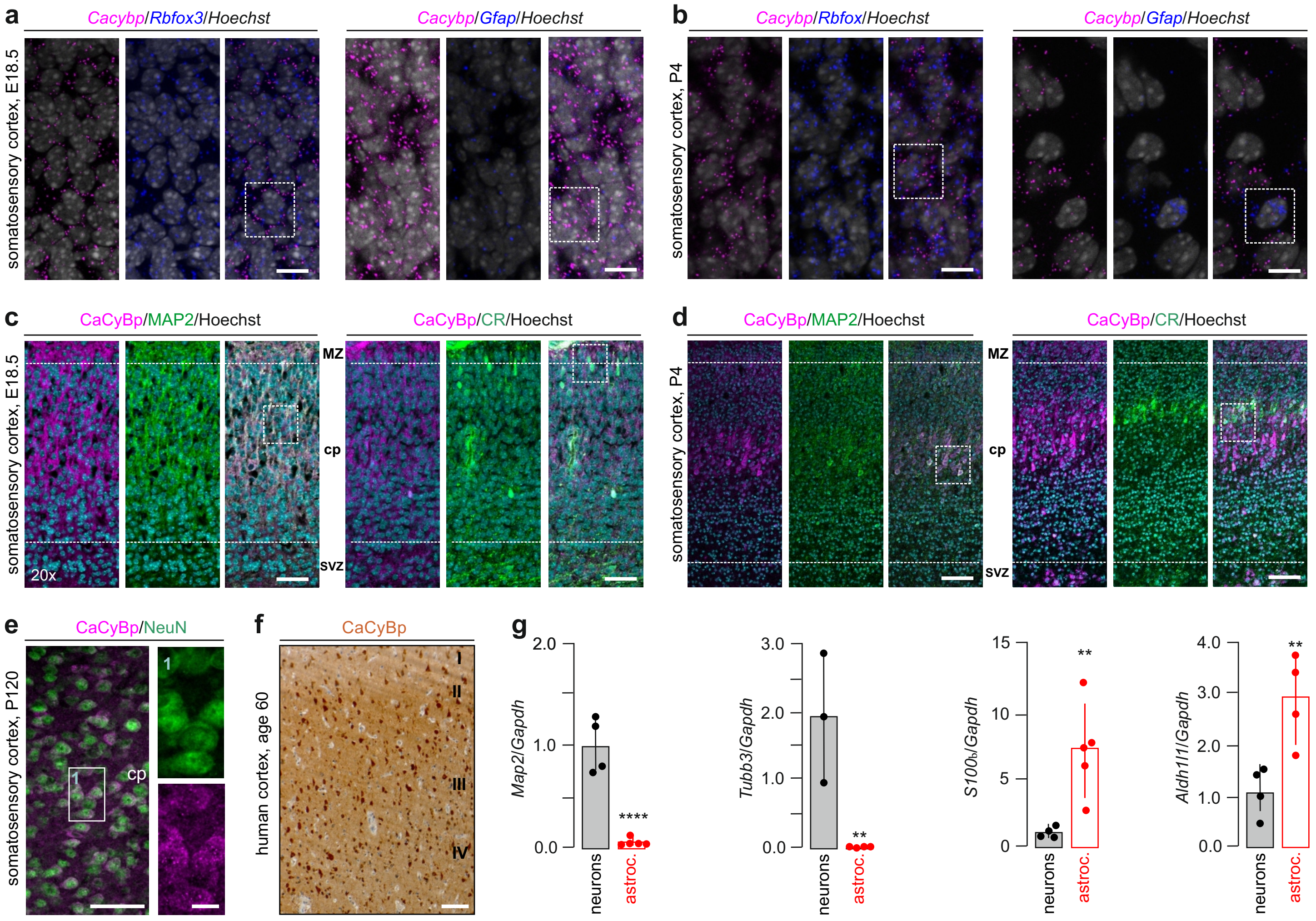

### Supplementary Figure 4

Cinquina *et al.* - Supplementary Fig. 4 (revision)

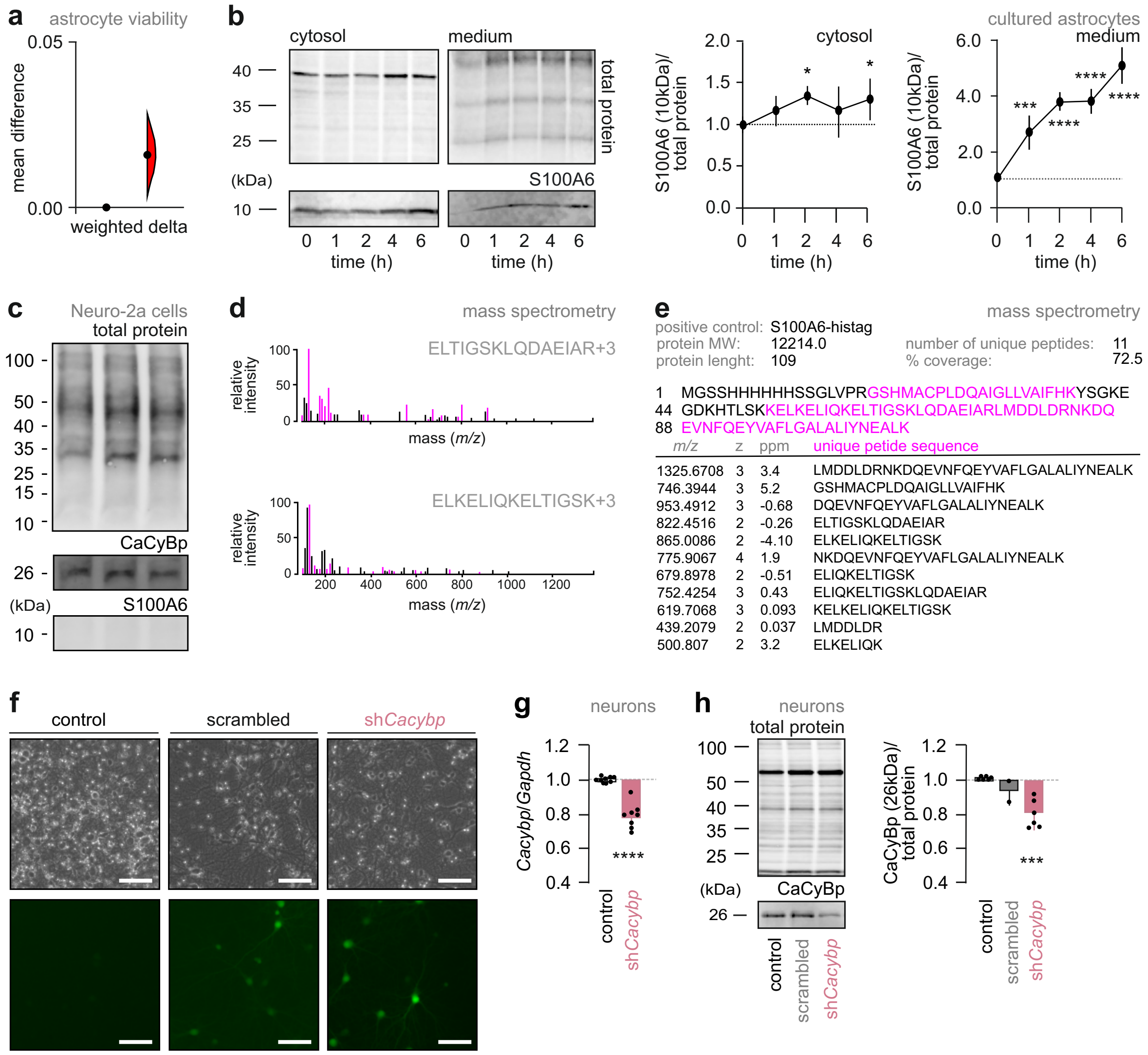

### Supplementary Figure 5

Cinquina *et al*. - Supplementary Fig. 5 (revision)

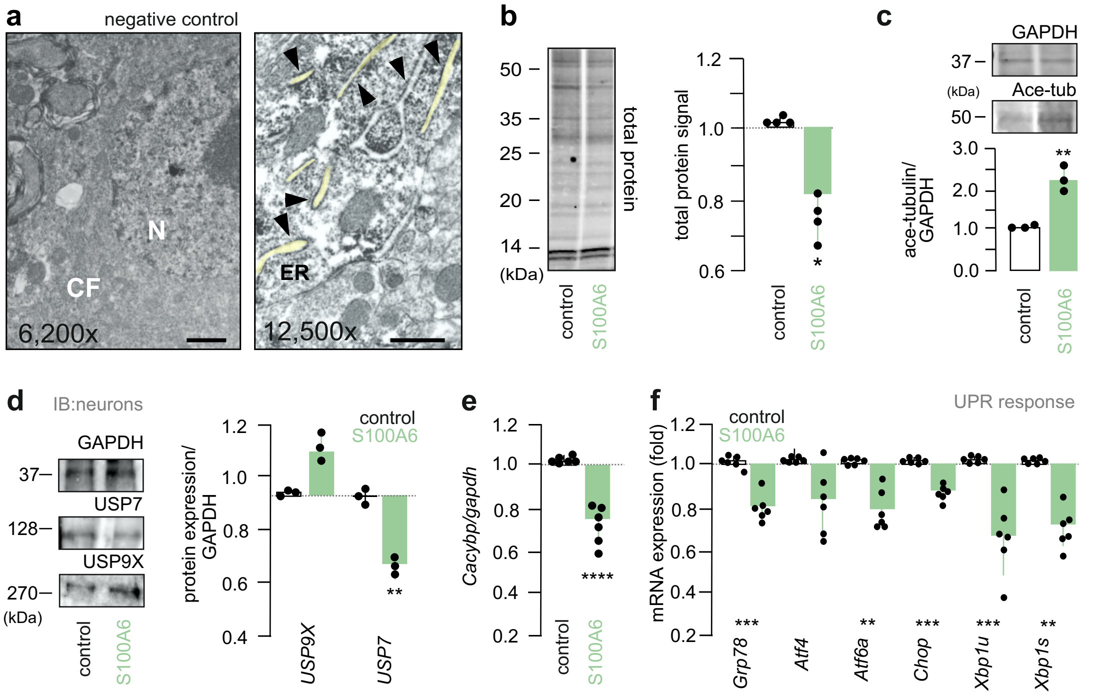

### Supplementary Figure 6

Cinquina *et al*. - Supplementary Fig. 6 (revision)

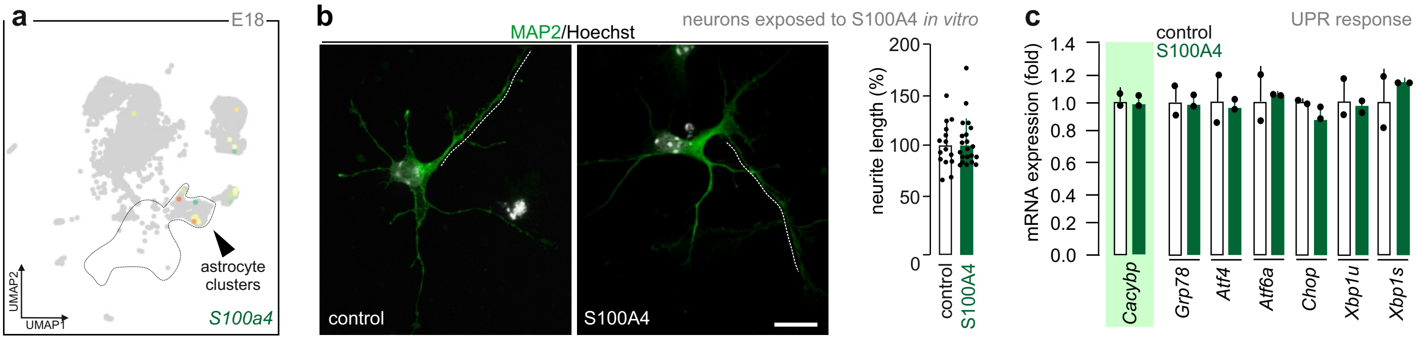

### Supplementary Figure 7

Cinquina *et al*. - Supplementary Fig. 7 (revision)

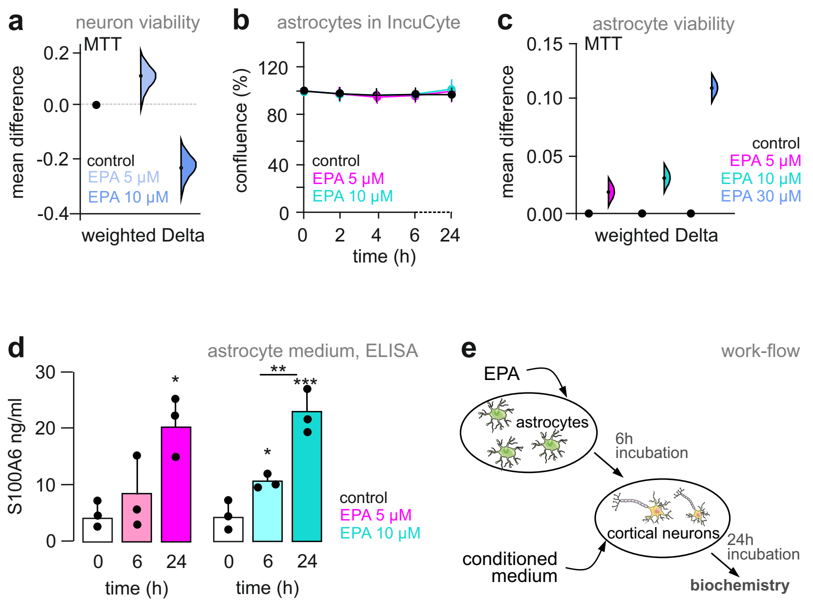

### Supplementary Figure 8

Cinquina *et al*. - Supplementary Fig. 8 (revision)

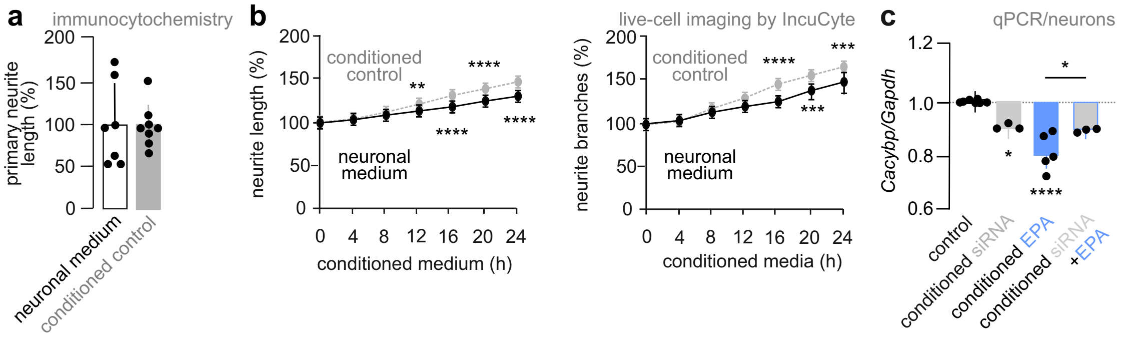

### Supplementary Figure 9

Cinquina *et al*. - Supplementary Fig. 9 (revision)

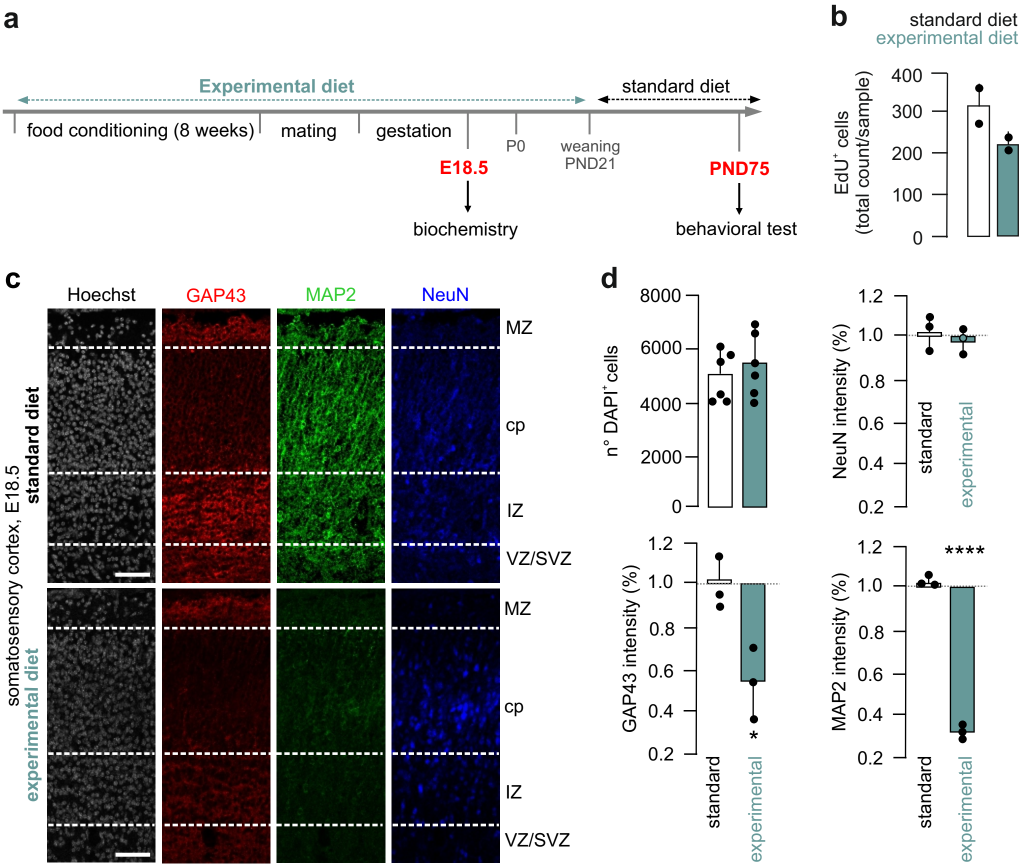
