## Supplementary Table 1 for "Astrocytes modulate neuronal development by S100A6 signaling"

Cinquina *et al*. - Supplementary Table 1 (revision)

| **Diet batch** | **Standard diet** | **Experimental diet** |
|---|---|---|
| Product name | 824050 | 823106 |
| Batch n. | | 37399 |
| date of manifacture | 09.02.05 | 18.08.14 |
| crude fat (CF) | 10.00% | 34.2% |
| Kcal/g fresh | 3.68% | 5.13% |
| Kcal/g 10% H20 | 3.47% | 4.81% |
| Vitamin E | 74.09 iu/Kg | 197.4% |
| C18:1 (n6)cis cis-12-Octadecanoic acid | n.r. | < 0.01% CF |
| C18:1 (n6)trans trans-12-Octadecanoic acid | n.r. | n.r. |
| C18:1 (n3)cis | n.r. | < 0.01% CF |
| C18:2 (n6)cis Linoleic acid (LA) | 1.34% | 10.5% CF |
| C18:2 (n6)trans Trans Linolelaidic acid | NA | 0.2% CF |
| **C18:3 (n3)cis Aplha-Linolenic acid (ALA)** | **0.23%** | **3.60% CF** |
| C18:3 (n6)cis Gamma Linoleic acid | n.r. | 0.1% CF |
| C18:4 (n3)cis Stearidonic acid | n.r. | 1.2% CF |
| C20:2(n6)cis cis-11,14-Eicosadienoic acid | n.r. | 0.8% CF |
| C20:3 (n3)cis cis-11,14,17-Eicosatrienoic acid | n.r. | 0.3% CF |
| C20:3 (n6)cis cis-8,11,14-Eicosatrienoic acid | n.r. | 0.2% CF |
| C20:4 (n3)cis cis-8,11,14,17-Eicosatetraenoic acid | n.r. | 0.8% CF |
| C20:4 (n6)cis Arachidonic acid (AA) | 0.01% | 0.4% CF |
| **C20:5 (n3)cis Eicosapentenoic acid (EPA)** | **n.r.** | **4.6% CF** |
| C22:2 (n6)cis Docosadienoic acid | n.r. | < 0.1% CF |
| C22:3 (n3)cis | n.r. | 0.1% CF |
| C22:4 (n6)cis Docosatetraenoic acid | n.r. | 0.1% CF |
| C22:5 (n6)cis cis-4,7,10,13,16 Docosapentaenoic acid | n.r. | 0.2% CF |
| C22:5 (n3)cis Docosapentaenoic acid (DPA) | n.r. | 1.7% CF |
| C22:6 (n3)cis Docosahexaenoic acid (DHA) | n.r. | 6.0% CF |
| C14:1 (n5) cis-9-Myristoleic acid | 0.01% | 0.1% CF |
| C16:1 (n7) cis-9-Palmitoleic acid | 0.07% | 3.7% CF |
| C18:1 (n9) cis Oleic acid | 0.7% | 9.13% CF |
| C12:0 (n3) cis Lauric acid | 0.07% | < 0.1% CF |
| C14:0 (n5) cis-9-Myristic acid | 0.13% | 3.3% CF |
| C16:0 (n7) cis-9-Palmitic acid | 0.23% | 11% CF |
| C18:0 (n6) Stearic acid | 0.14% | 2.7% CF |
| **Omega-3 fatty acids** | **0.26%** | **18.3% CF** |
| **Omega-6 fatty acids** | **1.87%** | **12.5% CF** |
| **omega-3/omega-6 ratio** | **1:7.2** | **1:0.7** |
