## Supplementary Table 2 for "Astrocytes modulate neuronal development by S100A6 signaling"

Cinquina *et al*. - Supplementary Table 2 (revision)

| | Gene name | experimental diet vs standard diet (fold) |
|---|---|---|
| **Proteosome / protein degradation/exosome** | *Ubap21* | 0,41 |
| | *Psme1* | 0,46 |
| | *Psmc6* | 0,48 |
| | *Rwdd1* | 0,60 |
| | *Serpina1* | 0,61 |
| | *Dnpep* | 0,71 |
| | *Ubap2* | 0,72 |
| | *Ncstn* | 0,79 |
| | *Ube213* | 1,34 |
| | *Ubxn1* | 1,36 |
| | *Npepps* | 1,44 |
| | *Psmd1* | 1,60 |
| | *Lonp1* | 1,61 |
| | *Ptub1* | 1,73 |
| | *Psmd6* | 2,07 |
| | *Psd11* | 2,56 |
| | *Psmc5* | 3,98 |

| | Gene name | experimental diet *vs* standard diet (fold) |
|---|---|---|
| **Cell proliferation/survival/apoptosis/cell cycle** | *Krt5* | 0,09 |
| | *Tppp3* | 0,26 |
| | *Manf* | 0,29 |
| | *Serbp1* | 0,36 |
| | *Lmna* | 0,39 |
| | *Caprin1* | 0,50 |
| | *Csrp2* | 0,56 |
| | *Erh* | 0,68 |
| | *Psmd13* | 1,25 |
| | *Fxr1* | 1,76 |
| | *Tsg101* | 1,80 |
| | *Srrt* | 2,00 |
| | *Ran* | 3,02 |
| | *Shc1* | 3,45 |

| | Gene name | experimental diet vs standard diet (fold) |
|---|---|---|
| **Translation / ribosome assembly / protein biogenesis /ER** | *Rplp2* | 0,22 |
| | *Pfdn2* | 0,26 |
| | *Rrbp1* | 0,27 |
| | *Pfdn6* | 0,29 |
| | *Rcn2* | 0,32 |
| | *Eef1b* | 0,38 |
| | *Rps25* | 0,40 |
| | *Fkbp3* | 0,47 |
| | *Rplp1* | 0,47 |
| | *Tma7* | 0,49 |
| | *Ssr1* | 0,50 |
| | *Eif4b* | 0,53 |
| | *Rpl23a* | 0,54 |
| | *Cnpy3* | 0,54 |
| | *Alad* | 0,55 |
| | *Tbca* | 0,58 |
| | *St13* | 0,59 |
| | *Sec23a* | 0,60 |
| | *Asna1* | 0,72 |
| | *Rpl24* | 1,42 |
| | *Eif2s1* | 1,42 |
| | *Rps26* | 1,46 |
| | *Rpl35* | 1,47 |
| | *Rpl22l1* | 1,47 |
| | *Kars* | 1,47 |
| | *Rpl7* | 1,49 |
| | *Nop56* | 1,58 |
| | *Rpl11* | 1,62 |
| | *Rpl13* | 1,69 |
| | *Gar1* | 1,71 |
| | *Rpl19* | 1,72 |
| | *Rpl17* | 1,73 |
| | *Rplp0* | 1,75 |
| | *Rps11* | 1,77 |
| | *Canx* | 1,78 |
| | *Fkbp10* | 1,78 |
| | *Rpl23* | 1,79 |
| | *Rpl6* | 1,88 |
| | *Ahsa1* | 1,90 |
| | *Hsph1* | 1,97 |
| | *Rpl7a* | 2,05 |
| | *Ppid* | 2,07 |
| | *Ganab* | 2,09 |
| | *Rpl8* | 2,20 |
| | *Rps2* | 2,30 |
| | *Rpl4* | 2,74 |
