## Supplementary Table 3 for "Astrocytes modulate neuronal development by S100A6 signaling"

Cinquina *et al.* - Supplementary Table 3 (revision)

| **Marker** | **Dilution** | **Host** | **Source** |
|---|---|---|---|
| CaCyBp | 1:1000 (IHC) | Rabbit | Human Protein Atlas/Sigma |
| Alexa Flour™ 546 Phalloidin | 1:500 | | Invitrogen |
| Beta III tubulin | 1:1000 | Mouse | Promega |
| Calnexin | 1:500 | Mouse | Invitrogen |
| Calretinin | 1:500 | Guinea Pig | Synaptic Systems |
| Cofilin | 1:500 | Rabbit | Cell Signaling |
| GAP43 | 1:500 | Rabbit | Millipore |
| GAPDH | 1:500 | Rabbit | Cell Signaling |
| MAP2 | 1:500 | Guinea Pig | Synaptic Systems |
| NeuN | 1:500 | Mouse | Merck/Chemicon |
| S100A6 (IHC) | 1:500 | Rabbit | Human Protein Atlas/Sigma |
| S100A6 (WB) | 1:500 | Rabbit | Aviva System Biology |
| USP7 | 1:500 | Rabbit | Bethyl Laboratories |
| USP9X | 1:500 | Rabbit | Bethyl Laboratories |
