## Supplementary Table 4 for "Astrocytes modulate neuronal development by S100A6 signaling"

Cinquina *et al*. - Supplementary Table 4

| Gene | | Primer sequence |
|---|---|---|
| *Actb* | (forward) | 5’-ATGGTGGGAATGGGTCAGAAG-3’ |
| | (reverse) | 5’-TCTCCATGTCGTCCCAGTTG-3’ |
| *Aldh1l1* | (forward) | 5’-AGCTGTGCCCTGAGTAATGT -3’ |
| | (reverse) | 5’-GCACAGCTTTGTTGAGGTCA -3’ |
| *Atf4* | (forward) | 5’-ATGGCCGGCTATGGATGAT-3’ |
| | (reverse) | 5’-CGAAGTCAAACTCTTTCAGATCCATT -3 |
| *Atf6a* | (forward) | 5’-GGACGAGGTGGTGTCAGAG -3’ |
| | (reverse) | 5’-GACAGCTCTTCGCTTTGGAC -3’ |
| *CaCyBp* | (forward) | 5’-GGTTGCTCCTCTTACAACAGG-3’ |
| | (reverse) | 5’-TGACCTCTCTGTGAAGTGCA-3’ |
| *Chop* | (forward) | 5’-CCAACAGAGGTCACACGCAC -3’ |
| | (reverse) | 5’-TGACTGGAATCTGGAGAGCGA -3’ |
| *Gapdh* | (forward) | 5’-AACTTTGGCATTGTGGAAGG -3’ |
| | (reverse) | 5’-ACACATTGGGGGTAGGAACA -3’ |
| *Grp78* | (forward) | 5’-ACCCTTACTCGGGCCAAATT-3’ |
| | (reverse) | 5’-AGAGCGGAACAGGTCCATGT-3’ |
| *Map2* | (forward) | 5’-CTTTCCCCTCTGGCTTCTGA -3’ |
| | (reverse) | 5’-AGCAAGGCATCTTCTCCACT -3’ |
| *S100a6* | (forward) | 5’-ACTCTGGCAAGGAAGGTGAC-3’ |
| | (reverse) | 5’-GCCGACATACTCCTGGAAGT-3’ |
| *S100b* | (forward) | 5’-TGTCTTCCACCAGTACTCCG -3’ |
| | (reverse) | 5’-TCCAGCGTCTCCATCACTTT -3’ |
| *Tubb3* | (forward) | 5’-CTCAACCACCTTGTGTCTGC -3’ |
| | (reverse) | 5’-GAAGAAGTGGAGACGTGGGA -3’ |
| *Xbp1s* | (forward) | 5’-GAGTCCGCAGCAGGTG -3’ |
| | (reverse) | 5’-GTGTCAGAGTCCATGGGA -3’ |
| *Xbp1u* | (forward) | 5’-GACAGAGAGTCAAACTAACTGGG -3’ |
| | (reverse) | 5’-GTCCAGCAGGCAAGAAGGT -3’ |
